# Topological data analysis to uncover the shape of immune responses during co-infection

**DOI:** 10.1101/723957

**Authors:** Karin Sasaki, Dunja Bruder, Esteban Hernandez-Vargas

## Abstract

Co-infections by multiple pathogens have important implications in many aspects of health, epidemiology and evolution. However, how to disentangle the contributing factors of the immune response when two infections take place at the same time is largely unexplored. Using data sets of the immune response during influenza-pneumococcal co-infection in mice, we employ here topological data analysis to simplify and visualise high dimensional data sets.

We identified persistent shapes of the simplicial complexes of the data in the three infection scenarios: single viral infection, single bacterial infection, and co-infection. The immune response was found to be distinct for each of the infection scenarios and we uncovered that the immune response during the co-infection has three phases and two transition points. During the first phase, its dynamics is inherited from its response to the primary (viral) infection. The immune response has an early (few hours post co-infection) and then modulates its response to finally react against the secondary (bacterial) infection. Between 18 to 26 hours post co-infection the nature of the immune response changes again and does no longer resembles either of the single infection scenarios.

**Author summary:** The mapper algorithm is a topological data analysis technique used for the qualitative analysis, simplification and visualisation of high dimensional data sets. It generates a low-dimensional image that captures topological and geometric information of the data set in high dimensional space, which can highlight groups of data points of interest and can guide further analysis and quantification.

To understand how the immune system evolves during the co-infection between viruses and bacteria, and the role of specific cytokines as contributing factors for these severe infections, we use Topological Data Analysis (TDA) along with an extensive semi-unsupervised parameter value grid search, and k-nearest neighbour analysis.

We find persistent shapes of the data in the three infection scenarios, single viral and bacterial infections and co-infection. The immune response is shown to be distinct for each of the infections scenarios and we uncover that the immune response during the co-infection has three phases and two transition points, a previously unknown property regarding the dynamics of the immune response during co-infection.

## Introduction

Co-infection is the simultaneous infection of a host by two or more phathogens. We are continuously exposed to multiple potential pathogens; many people are chronically (e.g. HIV) or latently (e.g. herpes viruses) infected, and we all carry potential pathogens in our colonising microbial flora. This means that nearly every new infection is some sort of co-infection, and globally, co-infections are the norm rather than the exception [1].

There is an impressive number of combinations of pathogens that derive synergy from contemporaneous infection of a host. These include viral-bacterial (e.g. influenza with pneumococcus or HIV with Mycobacterium tuberculosis), viral-viral (e.g. Hepatitis B with Hepatitis C), bacterial-bacterial (e.g. Borrelia burgdorferi with Anaplasma phagocytophila in Tick-borne illnesses) and pathogen-pathogen (e.g. Malaria with Dengue, Chikungunya, Filaria or Helminth) co-infections, to name a few. Although most studies to date have been focused on co-infections between two pathogens, infections with multiple pathogens are now becoming active topics of research [2].

Co-infections have effects on health at multiple levels: Co-infections can increase or decrease the rate of transmission of other infections [3], modulate the host immune response [4], create protection and resilience or susceptibility to further infections [5, 6], alter the performance of diagnostic tests and antimicrobial chemotherapy [7, 8], and even create opportunities for the emergence of new pathogens [9, 10]. In other words, some co-infections can have detrimental or even beneficial, outcomes.

The harmful effects of chronic co-infections, such as tuberculosis or Hepatitis B and C in association with HIV for example, are well established. However, generally and especially in acute infections, the mechanisms of co-pathogens with the host immune system and the possible consequences, ranging from insignificant, harmful or beneficial, are still largely unknown and difficult to dissect. The development of mathematical approaches that characterise the immune responses in the host have offered important steps for studying pathogen - pathogen interactions [11]. Interpretation of data sets with modern mathematical and machine-learning strategies can provide a comprehensive understanding of co-infections and their relevance/significance.

Topological Data Analysis (TDA) is a collection of computational tools derived from the mathematical subject of Algebraic Topology, that can help in identifying the behaviour of a biological system from a global perspective, guide detailed quantitative investigations and aid tailor further experimental settings. In fact, algorithms from topological data analysis have started to play important roles in novel interdisciplinary fields in biomedical sciences, including cancer genomics [12], diabetes [13], neuroscience [14], infectious diseases [15, 16], and in biology in general [17, 18].

Among the different TDA techniques for the qualitative analysis, the mapper algorithm [19] has shown a potential to simplify and visualize of high dimensional data sets. It generates a simple description of the data in form of a combinatorial object called a *simplicial complex*, that captures topological and geometric information of the point cloud in high dimensional space. The algorithm uses a (combination of) function(s) that map the data to a metric space, and builds an informative representation based on the clustering of subsets (which are associated to the values of the function(s)) of the data set. In the simplest case, this method reduces high dimensional data sets to a network whose nodes correspond to clusters in the data and edges to the existence of points in common between clusters. The aim of this algorithm is not to obtain a fully accurate representation of a data set, but rather a low-dimensional image which can highlight areas of interest, possibly for further analysis and quantification.

Using the mapper algorithm and data of malaria infection in mice, it was shown that the global shape of the stages a host infected with malaria goes through is circular; indicating a natural path infected individuals go through as they travel from health to sickness and back to recovery or death [15]. In [20] the authors developed a novel analytical tool based on persistent homology that helps to describe the geometric structure of the airways inside the lungs and can help in creating a more detailed classification of chronic obstructive pulmonary disease stages.

Motivated by the obvious potential of topological investigations in biomedical sciences, in the present study we seek to understand the evolution of the immune system as it responds to co-infection between virus and bacteria. Mathematical modeling research in influenza-pneumococcal co-infections has been a growing field within last years [4, 21–24]. These previous approaches are based on differential equations constructed based on biological reasoning. While they are suitable tools to test different hypothesis, these models are susceptible to bias by the designer and model complexity rapidly limits the reliability in the parameter fitting procedures [25].

Here, we use the co-infection data sets from [4] where we investigated the hierarchical effects of pro-inflammatory cytokines on the post-influenza susceptibility to pneumococcal co-infection by assessing the early and late kinetics of pro-inflammatory cytokines in the respiratory tract. In the experimental part of this study mice were divided into three groups and given either a single viral infection (with IAV strain A/PR8/34), a single bacterial infection (S. pneumoniae strain T4) or a co-infection (IAV + T4). The experimental readouts were the bacterial burden, viral titers and cytokine concentrations in the lung. Our previous work [4] used mathematical modelling that suggested a detrimental role of IFN-*γ* alone and in synergism with IL-6 and TNF-*α* in impaired bacterial clearance. We now use the mapper algorithm to investigate the global shape of the immune response under the three above infection scenarios, illustrated in Fig. 1.

**Fig 1.**
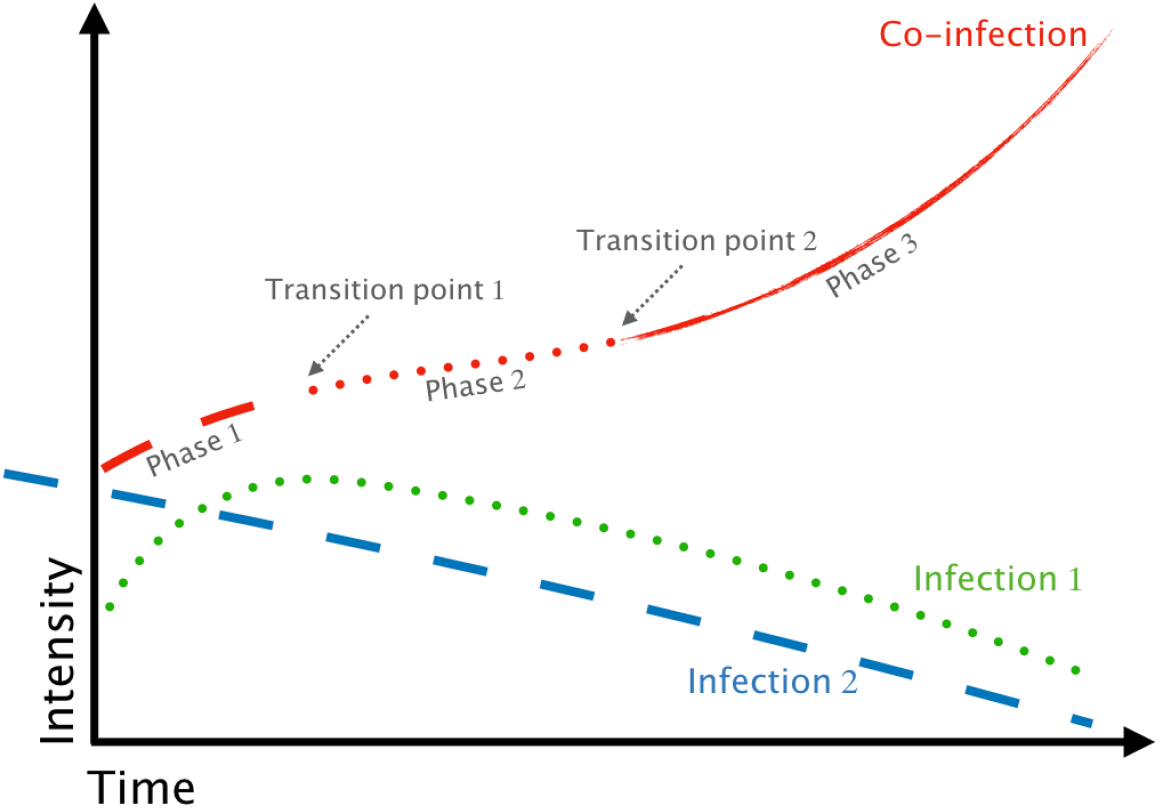
Predictions about the behaviour of the immune system in response to co-pathogenesis in the lung. Topological data analysis and nearest neighbour analysis reveal that initially the immune system inherits its behaviour from its response to the primary infection; it goes through a swift transition early in the co-infection (i.e. soon after the onset of the secondary infection) and it is consequently and temporarily driven mainly by its response to the secondary infection; There is a second transition point in the behaviour of the immune system and from there, it no longer resembles a standard response associated to either of the two single infections but rather it shapes its behaviour to respond to the co-infection itself.

## Results

### Persistent shape of the data in the three infection groups

We used the mapper algorithm to study four data sets. To start with, we gathered together the data for the three infection groups. The Kepler Mapper, a library implementing the mapper algorithm in Python, was employed [26]. Additionally, we wrote a semi-unsupervised algorithm that built all simplicial complexes for various metrics, lenses and ranges of values for the lens intervals and percentage overlap, and which chose simplicial complexes to represent the data. More specifically, using our semi-unsupervised algorithm, we tested the cosine, euclidean and correlation metrics, along with different epsilon values for the clusterer. Furthermore, all analysis was done using two lenses and we tested different combinations of the following: projections to the features of the data sets (i.e. the values of the pathogen load or the concentrations of the cytokines), the distance to the two nearest neighbours, the first two dimensions of various linear and non-linear dimensionality reduction algorithms and projections to (the image of) functions that reveal interesting geometric and statistical information about the data, such as density, eccentricity or centrality. We tested between 2 and 30 intervals for each lens and 10 different values for the percentage overlap of the lens’ intervals and the epsilon parameter of the clusterer. This resulted in almost 1 billion simplicial complexes being generated, from which, using properties of graphs such as the number of connected components, the algorithm chose complexes that were persistent in shape and that showcased important information about the immune response.

A detailed presentation of our semi-unsupervised algorithm and a discussion on how the representative simplicial complexes are chosen and how we came to the conclusion that those (simplicial complexes) are persistent in their shape is included in supplemental material S1. Of note during the parameter value search we obtained consistent results to those presented here with the same lenses and metric (correlation) and different parameter values for the number of intervals and percentage overlap. We also obtained similar results with the same metric but other pairs of lenses that included also linear and non-linear dimensionality reduction algorithms. Finally, we obtained similar results also with the cosine metric.

For the sake of clarity, we discuss only the results obtained by using the correlation metric with the following two types of lenses: lens 1 is the distance to the first neighbour and lens 2 is a projection to one of the features. Figs 2 and 3 illustrate the representative simplicial complexes that we discuss in more detail here (Table 1 in supplemental material S1 lists the parameter values specific for these simplicial complexes). Fig 2 corresponds to the data set that consists of all infection groups together, and Fig 3 shows the simplicial complexes for the individual infection groups separately. The columns in both figures indicate the projection for lens 2.

**Table 1.** Statistical study to data sets belonging to consecutive time points (columns) for each feature (row).

| Infection group | Feature | 1.5 - 6 | 6 - 18 | 18 - 26 | 26 - 31 |
| --- | --- | --- | --- | --- | --- |
| IAV | viral load lung | 0.912720 | 0.336585 | 0.607451 | 0.124077 |
| | IFN-γ | 0.925580 | 0.425324 | 0.685015 | 0.233908 |
| | TNF-α | 0.521103 | 0.673792 | 0.428918 | 0.489646 |
|  | MCP-1 | 0.951823 | 0.747659 | 0.812167 | 0.241850 |
|  | IL-6 | 0.725286 | 0.017479* | 0.143132 | 0.097698 |
| T4 | bacterial burden lung | 0.256184 | 0.409712 | 0.484568 | 0.683890 |
| | IFN-γ | 0.077142 | 0.186519 | 0.433551 | 0.963571 |
| | TNF-α | 0.382180 | 0.432187 | 0.012964** | 0.005679 |
|  | MCP-1 | 0.815624 | 0.618153 | 0.871879 | 0.900667 |
|  | IL-6 | 0.246575 | 0.634608 | 0.236886 | 0.928624 |
| IAV + T4 | Viral load lung | 0.113336 | 0.202011 | 0.000101*** | 0.200436 |
|  | Bacterial burden lung | 0.009209** | 0.006859** | 0.104765 | 0.958299 |
| | IFN-γ | 0.748814 | 0.269034 | 0.161643 | 0.320453 |
| | TNF-α | 0.520351 | 0.120874 | 0.016524* | 0.766774 |
|  | MCP-1 | 0.411175 | 0.353125 | 0.000691*** | 0.859898 |
|  | IL-6 | 0.879291 | 0.425412 | 0.019389* | 0.994039 |

p-values below 0.05 (\*), 0.01 (\*\*) and 0.005 (\*\*\*) are indicated.

**Fig 2.**
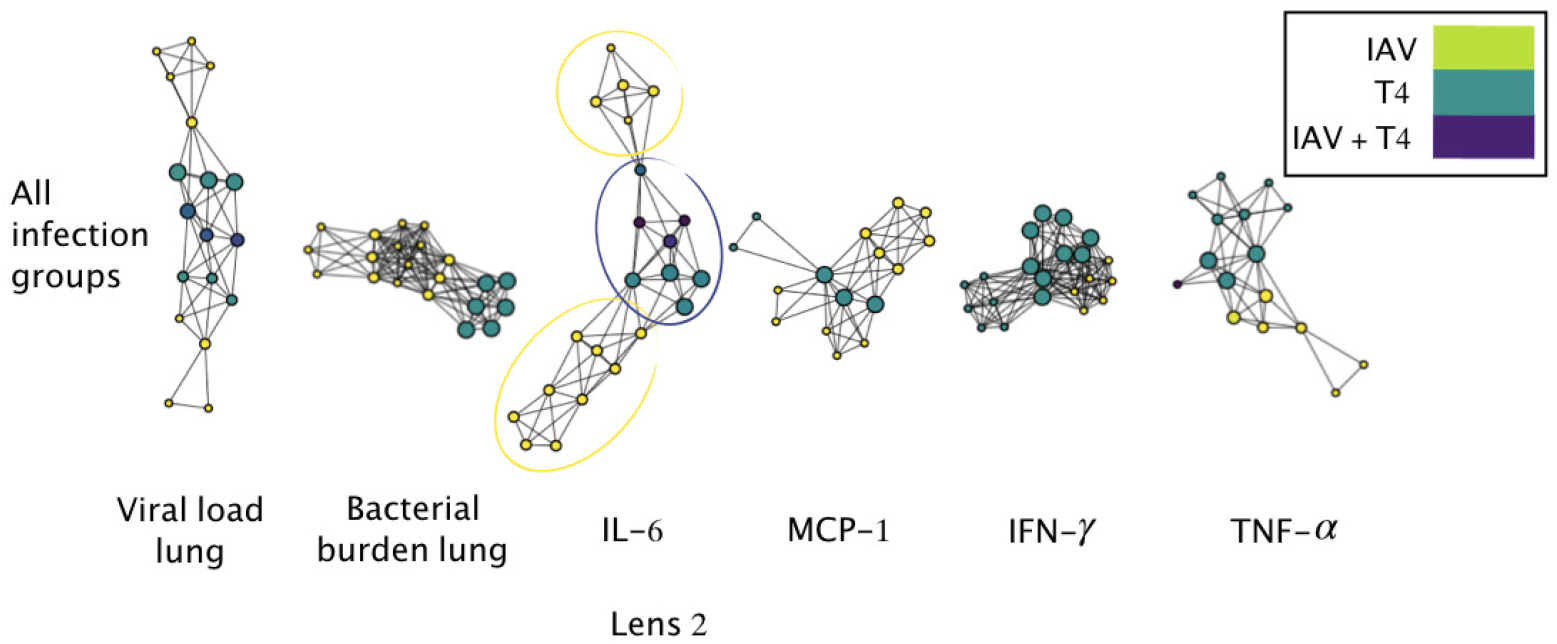
Simplicial complexes of data set consisting of all infection groups. The vertices of the simplicial complexes are color coded according to the infection group of the data points in the clusters. The legend shows which colors correspond to which infection group. Clusters that contain data points belonging to more than one infection group are colored by the average color value for members in that node.

**Fig 3.**
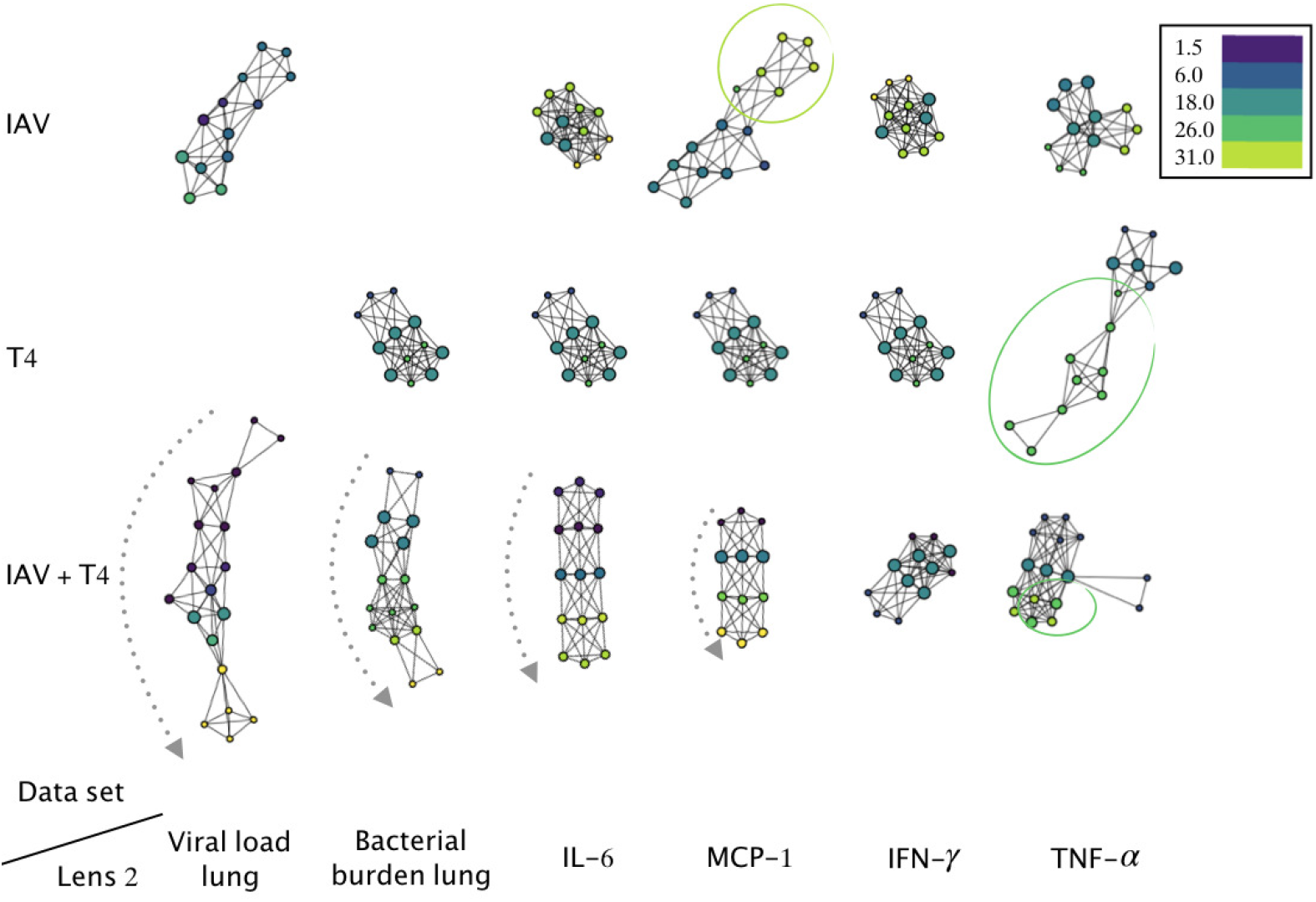
Decoupling simplicial complexes of the immune response to virus, bacteria and co-infection. Five simplicial complexes are generated for the single viral (top row) and single bacterial (middle row) infection groups and six simplicial complexes are generated for the co-infection group (bottom row) and. The vertices of the simplicial complexes are color (see legends) coded according to the hour post infection or co-infection in the clusters. Clusters that contain data subsets belonging to more than one time point are colored by the average color value for members in that node.

These analyses revealed persistence in the shape of the data. For example, in Fig 2 all the simplicial complexes generated with the different projections for lens 2 can be divided into three regions, two in yellow and one in purple/teal, where the circles (or vertices) in the yellow regions belong to time points 26 and 31 hours post co-infection (hpc) in the IAV + T4 infection group, and the circles in purple/teal belong to the IAV, T4 single-infection groups and early (1.5, 6, 18 hpc) time points in the IAV + T4 co-infection group. This is illustrated more explicitly in the simplicial complex generated by lens 2 = IL-6. The fact that the same shape is generated regardless of the projection used indicates that the shape is likely to represent the data.

Similar observations can be made for the simplicial complexes of the individual infection groups in Fig 3. For example, the simplicial complexes of the IAV infection group (second row) generated with the cytokines can all be divided into two regions, green vertices versus blue vertices. The simplicial complexes of the single bacterial infection (T4 infection group) generated with the bacterial burden and the cytokines IL-6, MCP-1 and IFN-*γ* are also all persistent in shape and, unlike the simplicial complexes for the other data sets, these ones do not highlight particular groups of nodes. The bottom row of Fig 3 corresponds to the simplicial complexes of the co-infection group (IAV + T4) alone. The time course can be clearly distinguished by the simplicial complexes generated with lens 2 being the two pathogens burden and the concentration of the cytokines IL-6 and MCP-1, as is indicated by the gray arrow (from early to later time points).

The simplicial complexes reveal different persistent shapes for the three infection groups - in IAV the later time points are segregated from the earlier time points; in T4 the simplicial complexes are homogeneous and do not reveal special areas; in IAV + T4 the time course of the infection is elucidated. Together this indicates that the immune system behaves differently in the three infection scenarios.

### Transition points in the immune response

We originally applied the mapper algorithm to all infection groups together in order to see whether the three infection groups would be clearly separated. However, the result illustrated in Fig 2 contains far more information. Taking the complex generated with the projection to IL-6 as the representative shape of the data, we observe that it has three regions: two in yellow and one in purple/teal. The yellow vertices of the simplicial complex correspond exclusively to all the late (26 and 31 hpc) time points of the co-infection group (IAV + T4), and the vertices in purple/teal correspond to the early (1.5, 6 and 18 hpc) data points of the co-infection group and to all the data points of the single viral (IAV) and single bacterial (T4) infection groups. As observed before, the simplicial complexes that are generated using the projection to the other features also highlight these regions. This indicates that in the later (26 and 31 hpc) time points of the co-infection the immune system behaves differently compared to its behavior in the single infection groups or at earlier time points during co-infection. This may imply a level of similarity in the behaviour of the immune system during the earlier time points in the co-infection and in the single infection groups. Together, from these two observations we can conclude that the topological data analysis has highlighted a transition in the nature of the immune response during a co-infection sometime between 18 and 26 hours post co-infection.

In order to interpret more precisely why the mapper algorithm is highlighting the specific regions illustrated inside the yellow circles, we calculated the p-values between groups of data points at consecutive time points, for each feature of the data (Table 1); we also made box plots to compare the data belonging to the two groups separated by the simplicial complex in Fig 2 corresponding to lens 2 = IL-6 (Fig 6 and Section 2.4 in S1 Text). The p-values show that, in the co-infection scenario, there is a strong change in the concentration of the cytokines TNF-*α*, MCP-1 and IL-6 (as well as in the viral load) between 18 and 26 hpc. The boxplots clearly revealed that data points of IAV + T4 from 26 and 31 hours post co-infection are separated by the simplicial complex from the earlier data points and that 18 and 26 hpc represent a transition point for the system, since at both time points there are data subsets that belong to both groups in the simplicial complex (the yellow and the teal/purple). It is possible that the shape of the simplicial complexes in Fig 2 is exactly mirrowing the dramatic change in the concentration of the cytokines (and possibly the viral load) that is revealed by the p-values.

Fig 4A shows the k-nearest neighbours for the data sets in the three infection groups, with metric correlation and 30 neighbours. The rows and columns indicate each data points and dark regions indicate the 30 closest neighbours of each data point. The labels show which points belong to which infection group and time point. The bottom right panel, inside the orange dotted box, corresponds specifically to the data points of the co-infection group (IAV + T4). The data sets in this region can be divided into three groups: the early period (1.5 hpc), the transition period (6 hpc) and the later period (between 18 and 31 hpc). This is illustrated by the fact that within the orange dotted box, the data sets at 1.5 hpc are neighbours with only data sets at 1.5 and 6.0 hpc, the data sets in 6 hcp have neighbours in both the early and the later groups and data sets at 18, 26 and 31 hpc have neighbours only at 6 hpc and late time points. This can be seen more clearly in Fig 4B which shows the correlation coefficients between data points in the co-infection data set. Data sets at 1.5 hpc are closely correlated to themselves, data sets at 6 hpc have close correlation to points in both the early and later periods, the later data sets (18, 26 and 31 hpc) are closely correlated to themselves only. Data sets between the early and later groups are not correlated.

**Fig 4.**
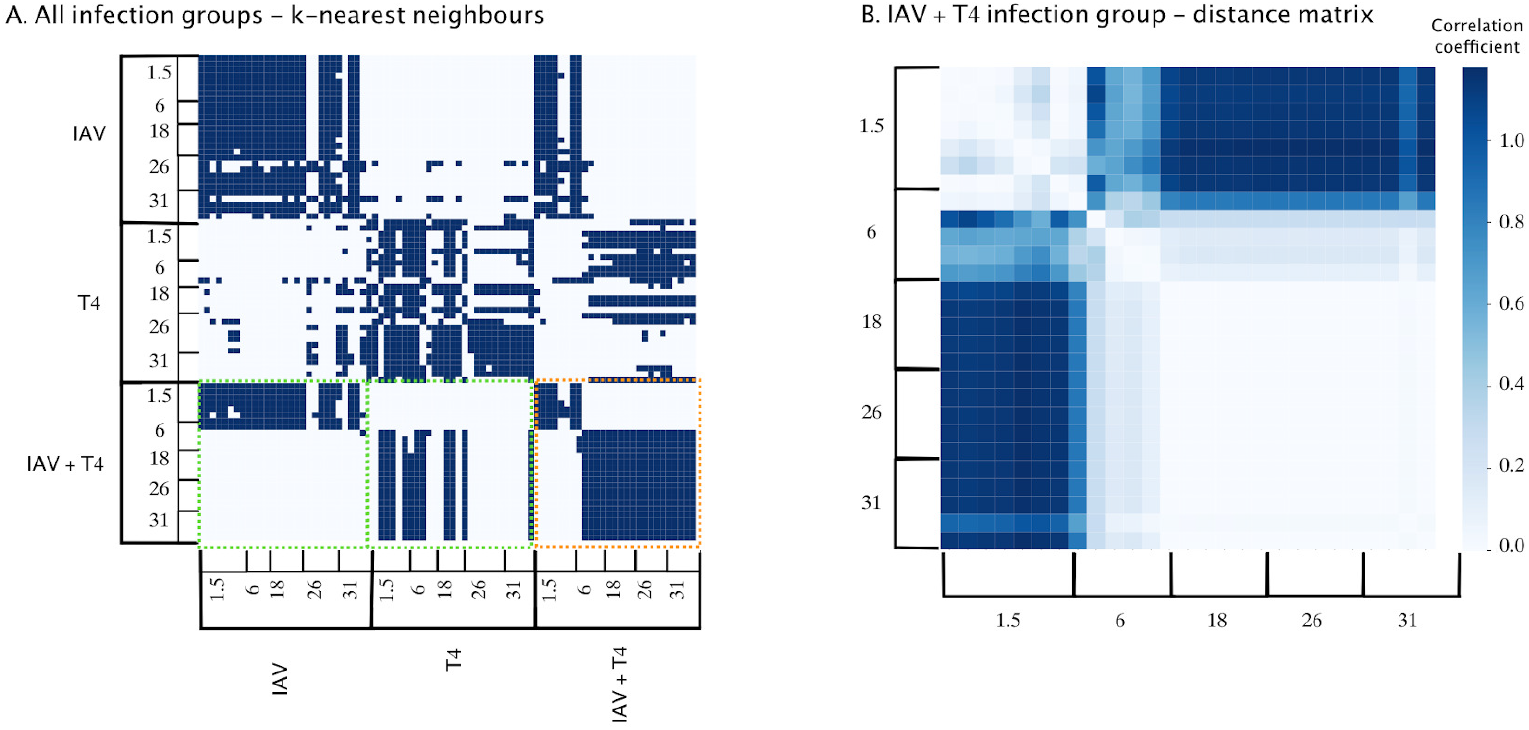
K-nearest neighbour analysis of co-infection. A) 30-nearest neighbours of all data sets in the three infection scenarios, with the correlation as metric. The infection groups and the time points for each infection group are indicated on the axes. The dark regions indicate the 30 neighbours of each data point. Inside the orange dotted box are the neighbours of co-infection data within the co-infection scenario. Inside the green dotted boxes are the neighbours of the co-infection data in the IAV single infection and T4 single infection groups. B) Correlation distance matrix of the co-infection group. The color-bar on the right indicates the correlation distance values between data sets. White indicates closely correlated and dark blue indicates not correlated.

In Fig 4A, now looking at the neighbours of the co-infection data subsets outside of the orange dotted box and inside the green dotted boxes (i.e. along the rows), we can see that co-infection data sets at early times also have neighbours exclusively in the IAV single infection group and co-infection data sets at late times also have neighbours exclusively in the T4 single infection group. In other words, the early co-infection data sets are closely correlated with the data sets in the IAV single infection group and the late co-infection data sets are closely correlated with the data sets in the T4 single infection group. Putting these three observations together, we can conclude that the immune response in the co-infection inherits its early response from the primary viral infection but at 6 hpc it undergoes a shift and quickly shapes its response to defend against the secondary bacterial infection.

In the simplicial complexes illustrated in Fig 3 in both the T4 single infection and co-infection (IAV + T4) groups the simplicial complexes generated with the TNF-*α* projection draw attention to the later time points in both infection scenarios. Specifically, for T4, the points correspond to time 26 hpc and in IAV + T4 the data sets correspond to times 26 and 31 hpc. Table 1 highlights that in both the single bacterial infection and in the co-infection, the concentration of the TNF-*α* cytokine changes significantly between 18 and 26 hours post the onset of the bacterial infection. Our previous analysis [4] of experimental results for TNF-*α* showed levels were increasingly and significantly elevated in co-infected mice from 18 hpi on. Therefore the simplicial complexes in Fig 3 of the single bacterial infection and the co-infection generated by the projection to TNF-*α* are exactly revealing this dramatic change in the concentration of this cytokine at this late stage in the course of the co-infection.

## Discussion

The relative contributions of the immune system during co-infections and how they can help in laying out the evolution of the immune system in response to co-infections are largely fragmented. The complexity of multi-pathogen infections makes detailed dissection of contributing mechanisms and stages of the immune response, which may be non-linear and occur on different time scales, challenging. Recently, in conjunction with experimental data, theoretical approaches have been able to uncover infection control mechanisms, establish regulatory feedback, connect mechanisms across time scales, and determine the processes that dictate different disease outcomes [27]. In this study we aimed at continuing this effort and we used TDA and data of co-infection experiments [4] to investigate how the immune system evolves between different infections.

Using the Mapper Algorithm (Figs 2 and 3) in combination with nearest neighbour analysis (Fig 4) we have shown that the immune response during influenza-pneumococcal co-infection consists of three stages (Fig 1): It is initially shaped by the inherited response to the primary influenza infection. We call this phase 1 of the immune response. Subsequently, the system undergoes an abrupt transition at 6 hours post onset of the secondary bacterial infection as it quickly modulates itself and starts responding predominantly to the bacterial infection; this is phase 2 of the immune response. There is a second transition stage between 18 and 26 hours post co-infection after which the immune response does no longer resemble the behaviour under a single viral or bacterial infection, but presumably shapes its response to the co-infection itself; this stage we call phase 3.

In [4], kinetics of bacterial growth and clearance in the respiratory tract and blood following IAV-*S. pneumoniae* co-infection revealed a turning-point between 6 and 18 hours post onset of the secondary bacterial infection. In this study we have narrowed down the time of the turning point specifically to 6 hours post co-infection.

Experimental results [4] for IFN-*γ* showed that the co-infection led to an increase as early as 1.5 hpc and 6 hpc compared to the single IAV infection. The levels of IFN-*γ* remained constant compared to the underlying IAV infection for the later time points, but a significant increase was observed when compared to the single T4 infection. Finally, overshooting concentrations of IL-6 in the co-infected mice were also detected experimentally at 26 hpc and 31 hpc compared to the single T4 infection. The chemokine MCP-1 was experimentally found to be significantly increased in the IAV+ T4 group compared to the single T4 infected group and marginally increased to the IAV only group at 26 hpi and 31 hpi.

The simplicial complexes illustrated in Fig 3 perfetly matches these observations. More specifically, the experimental observations made in [4] regarding the temporal changes in concentrations of the cytokines throughout the co-infection coincide with the special regions highlighted by the simplicial complexes. We observe that the simplicial complexes generated by the mapper algorithm with a projection to the features could be representing exactly those dramatic changes in the values of the features and that the algorithm is able to separate data points with high relative concentrations of cytokines away from other data points. In the simplicial complexes illustrated in Fig 3 in both the T4 and IAV + T4 infection groups the simplicial complexes generated with the TNF-*α* projection draw attention to the later time points in both infection scenarios. Specifically, for T4, the data sets correspond to time 26 and in IAV + T4 the data correspond to times 26 and 31. We could further interpret these results as further supporting evidence that the immune response at this stage of the co-infection is primarily responding in a way that is similar to its response in the single bacterial infection.

The simplicial complex of the co-infection generated with the projection to IFN-*γ* in Fig 3 segregates data points at early times 1.5 and 6 hpc (in purple) away from the clusters of the other data sets, resembling the experimental observations regarding the concentration of IFN-*γ* in the co-infection compared with the single viral and bacterial infections. The simplicial complexes of the co-infection scenario generated by the projections to IL-6 and MCP-1 (and to the viral load and bacterial burden) reproduced the timeline of the infection course. It is interesting to contemplate the possibility that the simplicial complexes of the co-infection imply that the cytokines IL-6 and MCP-1 play a consistent role through the whole infection course in the co-infection scenario.

Looking in more detail at the simplicial complex corresponding to the projection to IL-6 on the top row of Fig 2, we can see that the two groups of yellow circles are sprouting from different regions in the complex; one yellow group sprouts from vertices in purple and the other from vertices in teal. Recall that the vertices of the simplicial complex represent clusters of points of the data set and edges between vertices indicate that there are data in common between clusters. Therefore, after further analysis of the data sets that are in common between the purple and yellow vertices (clusters) and teal and yellow vertices, we could elucidate that, in fact, the purple vertices have data sets exclusively in the single viral infection at early time points and vertices in teal have all the data sets in the single bacterial infection and some data sets in the late stages of the single viral infection. In other words, the yellow region emanating from the purple vertices is connected exclusively to early time points in the single viral infection and the other yellow region to the single bacterial infection and late time points in the viral infection.

We may speculate the connections between the late-stage co-infection data and the early-stage single viral infection data are hinting at a rebound in viral titre after bacterial infection is established, which is a property of the co-infection that was been observed in other studies (for example) [28].

In [4] mathematical modelling, we proposed IFN-*γ* as a key and sufficient modulator in the impairment of bacterial clearance and other detrimental effects specifically for IL-6 and TNF-*α* in bacterial clearance. At the current stage of application of the mapper algorithm for this particular data set, we are not able to draw specific conclusions regarding causal roles of the cytokines in the co-infection. We can only allude to possible involvement of cytokines at specific stages in the infection course (as we have done, for example, with TNF-*α*, IL-6 and MCP-1.) In other words, TDA can dissect the potential of the different cytokines to represent the whole data set during co-infections, however, it can not point out or reject a key role of a specific cytokine for the susceptibility to bacterial co-infections.

The simplicial complexes generated for T4 single infection with projections to all the features show a homogenous structure where no specific group of data points are segregated or highlighted. We believe this is due to a trichotomy of pneumococcal outcomes discovered using stability and bifurcation analysis in [24]. Additionally, the immune response has been found to go through three stages during its response to single pneumococcal lung infections [29].

In [15] the simplicial complex of the stages a host infected with malaria goes through is circular and serves as a map of the loop an individual goes through on its way from health, through sickness and recovery and back to health. It is to be expected that a topological approach to study infectious diseases where hosts recover would also reveal circular topological simplicial complexes. This is not the case for the data set we have used. For the three infection groups, the data sets do not contain information for the full course of the infections. More specifically, the data set of the single IAV infected group is based on data starting from day 7 post the onset of the viral infection. The data set for the co-infected groups is incomplete towards the end of the infection course because for ethical reasons the mice that developed a high morbidity had to be euthanized before the bacterial infection is resolved naturally. Nevertheless, we found hints of looping behaviour in the co-infection, as discussed in supplemental material.

It has been difficult to uncover the implications in the biological context with certainty respect to the structures of the simplicial complexes. For example, we speculate that the segregation of specific data sets represents striking changes in feature values - i.e. changes in concentration of cytokine of pathogen load from one time point to the others. This seems indeed to be the case as explained earlier in this discussion. However, for example, striking changes in bacterial burden and viral load in the co-infection are shown in Table 1, where the bacterial burden changes significantly between 1.5 and 6 hpc and between 6 and 18 hpc and the viral load changes significantly between 18 and 26 hpc. However, it is not clear exactly how these changes are represented in the simplicial complexes of Fig 3. Therefore further quantification is required in order to understand in more detail the information that the choice of lenses provide in the biological context.

Mechanistic modeling studies investigating stages of the immune system in response to co-infections have been done previously [4, 21, 22, 29]. While under specific assumptions these models can highlight relevant mechanism, the design of mechanistic models and the abstraction complexity remain largely debatable. Here, TDA is presented as an additional tool to abstract high dimension data sets during co-infections, thereby significantly extending current knowledge and building a basis for translating improved mathematical models into potential therapies.

## Methods and Materials

### Experimental data

We consider the murine data that we first presented in [4]. In short, 7-8 weeks old C57BL/6J wild type mice were divided into three groups: single viral infection (IAV), single bacterial infection (T4) and co-infection (IAV + T4). The mice were anesthetized and intranasally infected with either a sublethal dose of IAV (A/PR8/34) or a bacterial infection with the *S. pneumoniae* strain T4 on day 7, or both, depending on the infection group. Following infection, mice were monitored daily for morbidity and mortality. Bronchoalveolar lavage (BAL), post-lavage lung and blood were collected at 1.5, 6, 18, 26 and 31 hours post bacterial infection (hpi) or post bacterial co-infection (hpc). Lungs were homogenized, and the supernatants were used to determine virus titers, immune cell populations, and cytokine and chemokine concentrations. Kinetic measurements for viral titers (mRNA by real-time PCR), bacterial counts (colony forming units (CFU)) as well as the following cytokines were considered: IFN-*γ*, TNF-*α*, IL-6, IFN-*β*, IL-22 and the chemokines MCP-1 and GM-CSF. The materials and methods are described in detail in [4].

For the analysis done in this study we took the measurements collected from the post-lavage lung. For the T4 and AIV + T4 infection groups, at 18 hpi and 18 hpc, respectively, the measurements had to be repeated. As it is a cross-sectional study (each measurement is coming from a mouse), we allocate high values of bacterial burden to rows that contain high values of cytokines. Mice were naive, we replaced N/A values (under level of detection) in the bacterial burden (in the single IAV infection group) and viral load (in the single T4 infection group) with the value 0. For each infection group, for time points that have less than five missing values in one feature, we replaced the missing values with the average value of the feature for that specific time point. For infection groups that have missing values at one single time point for one particular feature, we performed a linear interpolation between the mean values of the previous and the next time points and replaced the missing values with the predicted value for that time point.

### Data Analysis

Figs 2 and 3 were generated by performing topological data analysis (the Mapper Algorithm) on the single viral and bacterial infection and co-infection data sets using the Keppler-Mapper Python library [26]. Nodes in the simplicial complex represent clusters of infected mice, and edges connect nodes that contain samples in common. Nodes are colored by the average value of their samples for the variables listed in the Figs’ legends and color maps.

Three types of parameters are needed to generate a topological model: First is a notion of similarity, called a metric, which measures the distance between two points in some space (in our case, the points are the rows in the data and the space is a multidimensional space that can be plotted using the quantitative measurements of disease symptoms as axes, such as the pathogen load or cytokine concentration, i.e. the features of the data set). The metric we used is the correlation distance. Second are lenses, which are functions that describe the distribution of data in a space. A lens is a mathematical mapping (function) that converts a data set into a vector, in which each row in the original data set contributes to a real number in the vector; i.e. a lens operation turns every row into a single number. Metrics are used with lenses to construct the simplicial complex output. Multiple lenses can be used in each analysis. In this case, Keppler-Mapper handles them mathematically by considering the Cartesian product. Third is the resolution, which controls the number of bin partitions that will be created within the range of selected lens values, known as the number of intervals, and the amount of oversampling between bins, known as the percentage overlap. Clustering then takes place within the bins, forming the final vertices of the simplicial complex; and clusters are connected with an edge whenever they share data points within the region that are over-sampled according to the percentage overlap. Therefore increasing the number of bins increases the number of vertices and increasing the percentage overlap results in an increased number of edges. The metric, lenses, resolution, and clusterer used to generate the topological graphs in Figs 2 and 3 are as indicated in Table 1 in S1 Text.

K-Nearest Neighbour Analysis was implemented using the KNeighborsClassifier from the Scikit Learn Python library [30], with the correlation metric. T-tests were done using the ttest ind function from the SciPy Python library.

## Supporting information

### S1 Text. Supplementary Text

Detailed description of the computational methodology implemented in this study and presentation of further supporting material for the results and discussion sections. This includes the following supplementary figures and table: S1 Fig. 1, S1 Fig. 2, S1 Fig. 3, S1 Fig. 4, S1 Fig. 5, S1 Fig. 6, S1 Table. 1.

## Acknowledgments

We are grateful to the authors of the Keppler Mapper Python library who diligently answered our questions on the use of the library.

