## Supplemental for "Topological data analysis to uncover the shape of immune responses during co-infection"

### 1 Supplementary Material to Methods

#### 1.1 Topological Data Analysis

In this section we give a brief and intuitive introduction to the mapper algorithm and the algebraic topology concepts behind it. We also present in detail our methodology for using the mapper algorithm to investigate influenza in co-infection with bacteria. For further details, we refer the reader to the following resources: the original paper of the mapper algorithm is [1]. The paper presenting the Kepler-Mapper Python Library that we used to implement the mapper algorithm computationally is [2]. We recommend [3] as an introductory book to computational algebraic topology (albeit it does not include the mapper algorithm) and [4] for a thorough treatment of the mathematical subject of algebraic topology.

##### 1.1.1 Summary of steps for the implementation of the mapper algorithm

The mapper algorithm is a method of replacing a topological space by a simpler one, known as a *simplicial complex*, which captures topological and geometric features of the original space. The purpose of doing this is not only to obtain a visualization of high dimensional data sets in 3-d, but also because mathematical properties of simplicial complexes allow for the implementation of algebraic calculations that facilitate the classification of the topological features of the complex, and by extension, of the original topological space.

The algorithm begins with a data set of interest that consists of a point cloud $X$ containing $N$ points $x \in M$ sampled from a space $M$ whose topology we want to elucidate. We define a real valued function $f : X \to R$ (referred to in the literature as a *lens* or *filter*) whose value is known for the $N$ data points. Next, we find the range $I$ of the function $f$ that is restricted to the points in $X$. We divide this range into a set $S$ of smaller intervals of the same size, that overlap. This results in two parameters that can be used to control how detailed a representation of the data, i.e. the “resolution”, namely the number $l$ of the smaller intervals and the percentage overlap $q$ between successive intervals.

For each interval $I_j \in S$, we find the set $X_j = \{x|f(x) \in I_j\}$, i.e. the points in X that form its domain. For each set $X_j$ we form clusters $\{X_{jk}\}$, where $x_k \in X_j$ and $k >= 1$. We treat each cluster as a vertex in the resulting simplicial complex and draw

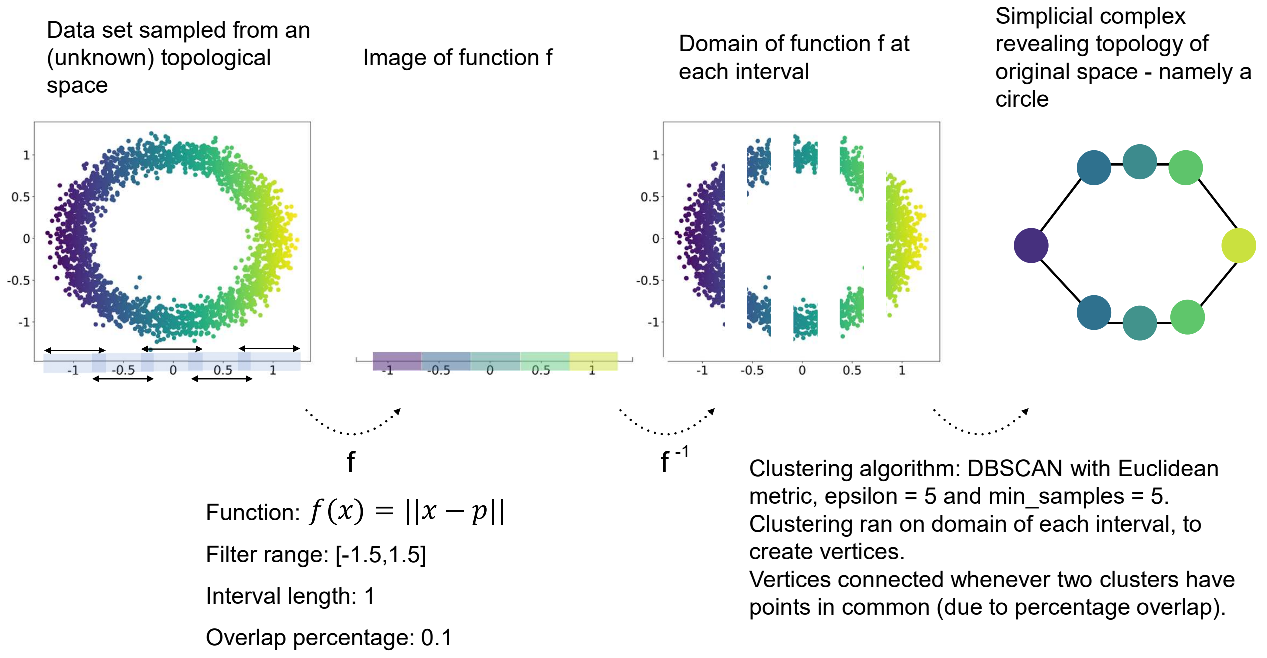


**S1 Fig. 1. Visualisation of the steps of the mapper algorithm being applied to a set of points sampled from the 2-dimensional circle.** Parameter $p$ in the function used as a lens is the leftmost point in the data. Example adapted from [1].

an edge between vertices whenever $X_{jk} \cap X_{lm} \neq \emptyset$, for $X_{jk} \neq X_{lm}$ (i.e. when different clusters have non-empty intersection). S1 Fig. 1 illustrates these steps for the construction of a topological network from data sampled from a 2-dimensional circle. The steps implemented computationally using the Kepler mapper python library [2].

##### 1.1.2 Simplicial complexes

To arrive at a precise definition of a simplicial complex and to understand it is not required for the context of this paper. Instead, here it is sufficient to think of simplicial complexes as a generalisation of networks that include higher dimensional elements. That is, simplicial complexes can be thought of as combinatorial objects consisting of vertices (0-d), edges (1-d), triangles (2-d), tetrahedra ("triangular pyramid") (3-d) and higher-dimensional ($n$-d for $n >= 0$) convex polyhedra. S1 Fig.2 illustrates this visually with a simple example.

In practical terms regarding the mapper algorithm, when we use one lens to construct a simplicial complex, the maximum dimension of the simplices is 1, i.e. the

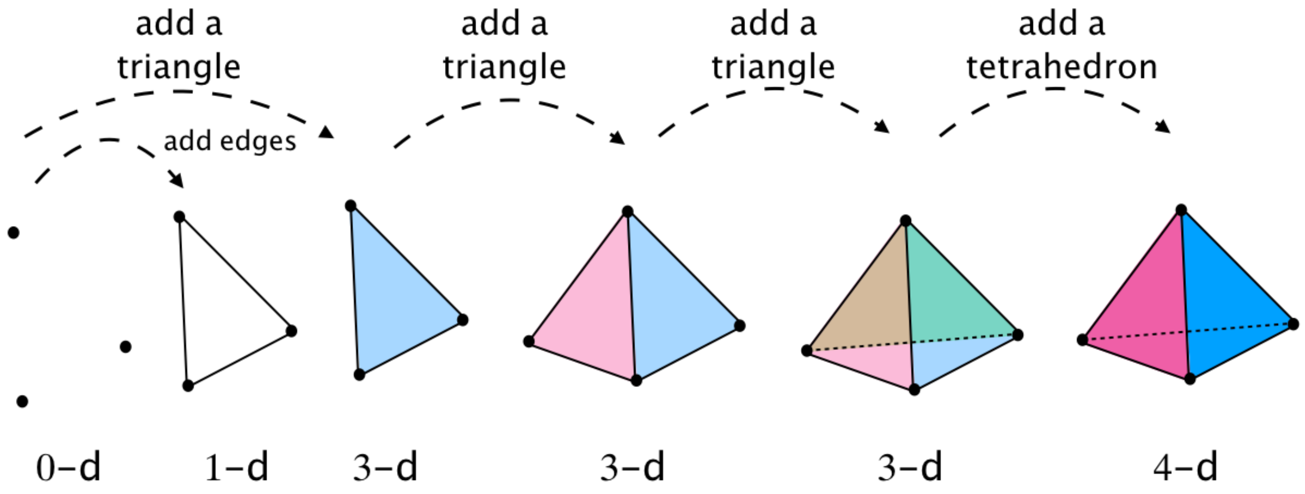


**S1 Fig. 2. A simple simplicial complex.** A simplicial complex can be thought of as a generalisation of a network, that also includes triangles, tetrahedra and higher dimensional convex polyhedra.

complex is made up of vertices and edges and is thus a standard network. When we use two lenses to construct a simplicial complex, the maximum possible dimension of the simplices is 4, meaning it is made of vertices, edges, triangles and tetrahedra. Section 3.2 in the original paper of the mapper algorithm [1] describes explicitly how simplicial complexes are built with 2 lenses and how this can be generalised to more lenses and higher dimensional simplicial complexes.

###### 1.1.3 Choosing the metric space(s)

The important aspect about the choice of metric for the purposes of the mapper algorithm is to have a notion of distance between two data points.

Although the mapper algorithm is less sensitive to the choice of metric than other methods that aim creating simpler representative objects of spaces in high dimensions (e.g. dimensionality reduction algorithms) [1] we still wanted to make sure that our investigation was not biased by the choice of metric. Therefore we tested three metrics that "appeared suitable" for our data sets. These were the Euclidean, cosine and correlation metrics. We chose the Euclidean metric as it is the most intuitive to work with. Next, we wanted to investigate whether looping is a recurrent property in our data sets. In [5], where looping was indeed found to be a motif in their data set (of an infection from which patients can recover, such as in our study), the cosine metric was used, so we also decided to test this metric. Finally, we regarded the correlation metric to make the most biological sense. So we also tested this one.

Specifically, the distance is defined as follows for the three metrics we work with:

**Euclidean:** if $x = (x_1, ..., x_n)$ and $y = (y_1, ..., y_n)$ are two points in Euclidean $n$-space, then the Euclidean distance between them is given by the Pythagorean formula:

$$d(x, y) = \sqrt{\sum_{i=1}^{n} (y_i - x_i)^2}$$

Computationally, this is done using the `sklearn.metrics.pairwise.euclidean_distances` function from Scikit-learn.

**Cosine:** if $x = (x_1, ..., x_n)$ and $y = (y_1, ..., y_n)$ are two points in Euclidean $n$-space, we use the following bespoke distance definition:

$$d(x, y) = \left| \frac{x.y}{||x||_2 * ||y||_2} - 1 \right|$$

Where $x.y$ is the dot product between $x$ and $y$ and $||u||_2$ is the L2-norm (Euclidean norm) of $u$. This definition of the is the absolute value of the normalised dot product of $x$ and $y$ minus 1. The reason we take this definition and, not simply the normalised dot product, is the following: The normalised dot product of any two points takes values between +1 and -1. We want to cluster points whose vectors from the origin are almost parallel and point almost in the same direction. Two such points have a normalised dot product value that is close to +1. However, DBSCAN clusters two points whenever they are a distance less than or equal to the `eps` parameter. (The `eps` parameter is the maximum distance between two samples for them to be considered as in the same neighbourhood). Therefore, what we need is a distance formula that outputs small values for points that are "close" to each other (i.e. whose vectors from the origin point in roughly the same direction). The above bespoke distance formula achieves this. In order to implement this computationally, we precompute the distance matrix of the data using the normalised dot product with the `sklearn.metrics.pairwise.cosine_similarity` function of sklearn, next, we preprocess the distance matrix such that it fits our bespoke distance criterion:

```
X_cosine_similarity = sklearn.metrics.pairwise.cosine_similarity(X)
X_dist = np.abs(X_cosine_similarity - 1)
```

Finally, we pass the precomputed distance matrix to the clusterer, setting the `metric` parameter of DBSCAN to be equal to 'precomputed'.

**Correlation:** if $x = (x_1, ..., x_n)$ and $y = (y_1, ..., y_n)$ are two points in Euclidean $n$-space, then the correlation distance between them is given by the following formula:

$$d(x, y) = 1 - \frac{(x - \bar{x}).(y - \bar{y})}{||(x - \bar{x})||_2||(y - \bar{y})_2||}$$

Where $u.v$ is the dot product between $u$ and $v$, $\bar{u}$ is the mean of the elements of $u$ and $||u||_2$ is the L2-norm (the Euclidean norm) of $u$. Computationally, this is done using the `sklearn.metrics.pairwise_distances` function from Scikit-learn, setting the parameter `metric` to 'correlation'.

In the main part of this study we only reported the results obtained with the correlation metric. We observed the same results with the correlation and cosine metrics and we could not make concrete observations about the result obtained with the Euclidean metric. The questions of why the cosine and correlation metrics create the same simplicial complexes and how they differ from those created with the Euclidean metric require further and were not done as part of this study.

##### 1.1.4 Selection of lenses

The outcome of mapper algorithm is highly dependent on the lens(es) chosen. For the purposes of our study, we categorised the lenses we used according to the information about the data they extract.

**Features.** These lenses are simply the values at each data point of a particular feature of interest. These were calculated using the `fit_transform` function of the Kepler mapper.

**Distances to closest neighbours.** These lenses report the distance of each data point to its $n$ closest neighbours, or the sum of the distances to the $n$ closest neighbours, under the metric of choice. These were calculated using the `sklearn.metrics.pairwise_distances` function from Scikit-learn, specifying the metric to be one of the three described above.

**Dimensionality Reduction.** These are projections of the data, usually, to the first (and possibly also the second) dimension(s) of various dimensionality reduction algorithms. These were calculated using the Manifold Learning algorithms from the Scikit-learn Python library and the `sklearn.decomposition.PCA` and `sklearn.decomposition.TruncatedSVD` functions of Scikit-learn. See section 1.1.5 for a brief description of the methods used here.

**Geometric properties.** These report geometric properties. Specifically, we tested:

- The density using the `sklearn.neighbors.KernelDensity` function with Gaussian kernel and calculated the bandwidth using Scott's Rule [6].
- The eccentricity which is defined as follows:
  Given $p$ with $1 <= p < +\infty$, define
  $$E_p(x) = \left( \frac{\sum_{y \in X} d(x,y)^p}{N} \right)^{\frac{1}{p}}$$
  where $x, y \in X$. (Recall that we denote the data set of $N$ points by $X$ and $d(x,y)$ denotes the distance between $x, y \in X$, which is dependent on the metric of choice.)
- The infinite centrality which is a generalisation of the eccentricity above, for when $p = +\infty$, then $E_\infty(x) = max_{x' \in X} d(x, x')$.

**Statistical properties.** These report on statistical properties about the data points, such as the sum of the values of all the features for each data point, the average value of the features for each data point, etc. These were calculated using the `fit_transform` function of the Kepler mapper.

##### 1.1.5 Short explanation of dimensionality reduction algorithms

Here we provide short explanations about each of the dimensionality reduction algorithms used, (some of which have been adapted from the documentation of Scikit-learn [7]).

Singular value decomposition (SVD) and Principal Component Analysis (PCA) are two methods used to perform linear dimensionality reduction of high-dimensional data set.

PCA reduces the data into linearly uncorrelated variables (called *principal components*) such that the first component accounts for as much of the variability in the data as possible, and each succeeding component in turn accounts for the next highest variance possible and is orthogonal to the preceding components. When used in dimensionality reduction, one can take the first few principal components as the new set of features of the data set. To implement this, we used `sklearn.decomposition.PCA`.

SVD is a matrix decomposition method for reducing a matrix $A$ to the product of its constituent parts ($A = U \cdot \Sigma \cdot V^T$, where $U, V$ are unitary matrices and $\Sigma$ is a rectangular diagonal matrix of singular values). When applied in dimensionality reduction, one can select the top $k$ largest singular values in $\Sigma$ and use the $k$ affiliated columns in $\Sigma$ and rows in $V^T$ to generate an approximation $B = U \cdot \Sigma_k \cdot V_k^T$ to $A$. We used the TruncatedSVD class of Scikit-learn to implement this.

LLE, Isomap, MDS and Spectral Embedding are algorithms that belong to a framework called Manifold Learning, which are an attempt to generalize linear frameworks like PCA, to be sensitive to non-linear structure in data. These approaches learn the high-dimensional structure of the data from the data itself, in an unsupervised manner, i.e. without the use of predetermined classifications.

The Isomap seeks a lower-dimensional embedding which maintains geodesic distances between all points. LLE seeks a lower-dimensional projection of the data which preserves distances within local neighborhoods, and it can be thought of as a series of local Principal Component Analyses which are globally compared to find the best non-linear embedding. For the Isomap and the LLE one needs to make a choice on the number of neighbors; since in our data set each time point should have between two and five data points corresponding to individually analyzed mice, we chose the value of three neighbors for these algorithms.

Multidimensional scaling (MDS) seeks a low-dimensional representation of the data in which either the distances between the two output points are set to be as close as possible to the similarity or dissimilarity of the original data (the metric approach) or the algorithm seeks a monotonic relationship between the distances in the embedded space and the similarities/dissimilarities of the original data (the non-metric approach).

For spectral embedding Laplacian Eigenmaps are implemented, which find a low dimensional representation of the data using a spectral decomposition of the graph Laplacian. The graph generated can be considered as a discrete approximation of the low dimensional manifold in the high dimensional space. Points close to each other on the manifold are mapped close to each other in the low dimensional space, preserving local distances.

###### 1.1.6 On the number of intervals and percentage overlap for each lens

The choice of number of intervals and percentage overlap defines how detailed or coarse a topological network representation of the data we want to create is, in other words, the "resolution" of the representation. The number of intervals can be any number starting from 1 and, as the name indicates, the percentage overlap can be anything in the range $[0.0, 1.0]$. Thus, choosing a higher number of intervals translates to increasing the number of vertices of the graph. Increasing the percentage overlap allows the algorithm to find more data points in common between two intervals, thus roughly translating to an increased possibility of connecting two vertices. (Note that increasing the number of lenses also increases the number of vertices, since it increases the number of intervals that the data is partitioned into). S1 Fig. 3 shows topological networks of data sampled from a 2-dimensional circle, at different resolutions.

###### 1.1.7 Scaling of the data

PCA gives different results when the scales of the features are different. Additionally, manifold learning methods are based on a nearest-neighbor search, therefore, such algorithm may perform poorly if the features of the data are on different scales. Therefore, we normalise the data before applying the dimensionality reduction algorithms. (Note that the clustering algorithm is run on the original data, which we do not normalise.)

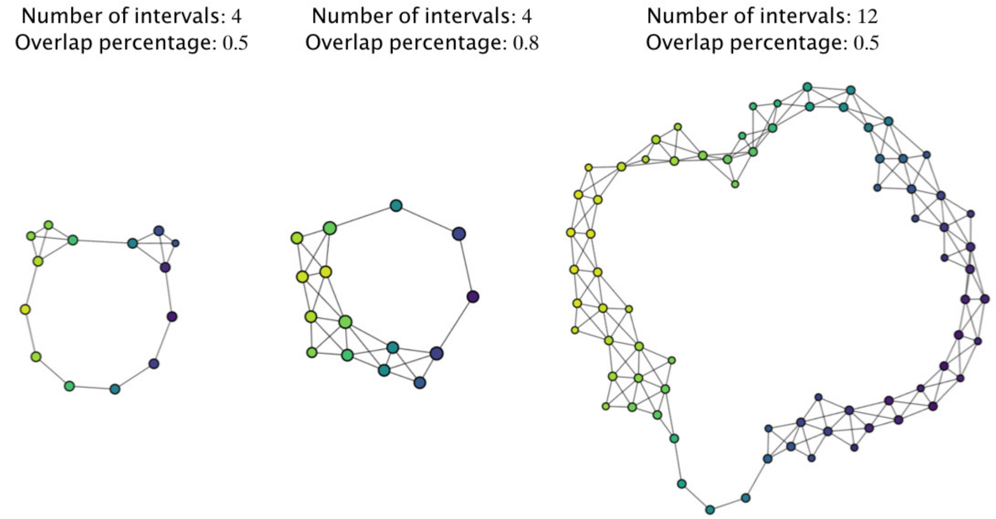


**S1 Fig. 3. Resolution of simplicial complexes and persistence of the shape of the data**. Three simplicial complexes of data sampled from an unknown 3-dimensional space of different resolution are generated using the mapper algorithm [1], implemented computationally using the Kepler mapper Python library [2]. One lens is used for all simplicial complexes, namely a projection on the x-axis. The number of intervals and percentage overlap for the lens at each resolution are indicated. Observation 1: the three simplicial complexes consist of one connected component. Observation 2: the three simplicial complexes have a "big hole" in the middle. Conclusion: The data set is sampled from a topological space that consists of one component and that has a hole in the middle. Since we know it is a 3-dimensional space, we can say it is a torus.

###### 1.1.8 Choice of clustering algorithm

Finding a good clustering of the points is a fundamental issue in computing representative topological networks. Currently there is no automated or principled method of making a choice. The mapper algorithm does not have any limitations on the cluster used and in particular, according to [1] desired characteristics when choosing a clustering are that it:

1. is able to take an interpoint distance matrix as an input, not restricted to the Euclidean distance,
2. does not require specifying the number of clusters.

We chose the Density-Based Spatial Clustering of Applications with Noise (DBSCAN) [7] because we can specify what metric to use, the thresholds for the number of points needed to create a cluster (`min_samples`) and the value of the distance below which two points are considered to belong to the same cluster (`eps`).

##### 1.2 Grid search analysis of parameter values for the mapper algorithm

As described in the previous section, the outcome of the mapper algorithm is highly dependent on the choice of values for the following parameters: the number of lenses, the type of lenses, the number of intervals for each lens, the percentage overlap for the intervals, the choice of metric space and the clustering algorithm.

We wrote a semi-unsupervised algorithm that built all simplicial complexes for various metrics, lenses and ranges of values for the lens intervals, percentage overlap and epsilon values of the DBSCAN clusterer from sklearn, and which "chose appropriate" simplicial complexes to represent the data. More specifically, below we list all the values we tested.

**Metrics:** cosine, euclidean and correlation

**Number of lenses:** 2

To start with, we constructed simplicial complexes of the data sets with between 1 and 4 lenses. However, when we visually inspected the result of using a three or four lenses, we observed many more clusters corresponding to the same points, because of oversampling by the larger number of intervals introduced by the third and fourth lenses. In addition, using more that two lenses can make the calculations take over 20 hours to complete, compared to 3-5 hours for two lenses (ran on a standard laptop). Therefore we limited ourselves to using two lenses.

**Combinations of lenses:**

1. Lens 1 = Distance to the first neighbour with Lens 2 = Distance to the second neighbour
2. Lens 1 = Distance to the first neighbour with Lens 2 = Projections to features
3. Lens 1 = Sum of the distances to the first and second neighbours with Lens 2 = Projections to features
4. Lens 1 = First dimension of a dimensionality reduction algorithm with Lens 2 = Projections to features
5. Lens 1 = First dimension of a dimensionality reduction algorithm with Lens 2 = Second dimension of the same dimensionality reduction algorithm
6. Lens 1 = Geometric or statistical information with Lens2 = Projections to features

The projections to features lenses are a projection to each of the nine features of the data sets (the two pathogen loads and the seven cytokine concentrations). The distance to the closest neighbours and the geometric and statistical lenses depend on the metric chosen .

**Number of intervals:** between 2 and 30.

**Percentage overlap:** between 0.1 and 0.9.

**Clusterer:** DBSCAN from scikit-learn [7]. For the `min_samples` parameter we chosed the value of 1, i.e. a cluster can be formed with 1 or more data points. For the `eps` parameter we tested values between the minimum and maximum distances in the data sets for each metric. More specifically, the distances between the data points in the three infection groups range between [0,520002] in the Euclidean metric; between [0,1] in the cosine metric; and between [0,2] in the correlation metric. So for the Euclidean metric we tested 5000 values between 0 and 520002; and for the cosine and correlation metrics we tested 10 values between 0 and 1 and 0 and 2, respectively.

Taking all these together, there are 18 different pairs of lenses; 28 different intervals and 10 percentage overlap values to test for each lens; 10 values for the epsilon parameter for the clusterer for the cosine and correlation metric and 5000 for the Euclidean metric. Therefore, in total this parameter grid search exercise generated almost 1 billion simplicial complexes $(15 * 28^2 * 9^2 * 10 + 3 * 28^2 * 10^2 * 5000 = 962,085,600)$.

From all the simplicial complexes generated, the algorithm chooses those that have a user-specified number of connected components. Biologically it makes sense that the simplicial complexes for the three infection groups have only one connected component. From the resulting list of simplicial complexes with one connected component, we ordered the simplexes in ascending order of epsilon value for the clusterer and the percentage overlap for the intervals for both lenses; choosing the smaller rather than the larger values for these parameters makes the simplicial complexes, as models of the system, simpler, which is desirable when working with models. Next, we chose simplicial complexes that had number of vertices at least half of the number of data points in the data set (for example, if the data set for the IAV infection group has 30 data points then we favour simplicial complexes that consist of only one connected component and at least 15 vertices); choosing this number gives a good resolution, not too simple, but also prevents the user from choosing simplicial complexes that have oversampling of data points. Finally, we visually inspected the simplest (in terms of small values for the parameters) 10, 20 or 30 simplicial complexes in that list. The visual inspection has two purposes: First, to give the user an idea of whether there is persistence in the global and local structures of the simplicial complexes generated. Second, to choose a representative structure for the data set.

### 2 Supplementary material to results/discussion

#### 2.1 Search for looping behaviour in co-infection

**Definition:** *Disease space* is the multidimensional space that can be plotted using quantitative measurements of disease symptoms as axes.

**Definition:** Parameters of a data set that oscillate, partially overlap and have a time lag between them are called *hysterectic parameters* [5].

**Definition:** When pairs of hysterectic parameters are plotted against each other in their phase plot they create loops [5]. These can be considered two dimensional projections of the higher dimensional looping behaviour of the data set and are referred to as *disease maps.*

In the context of infectious diseases from which hosts can recover (e.g. malaria, influenza, pneumococci, etc), disease maps that form loops are representative of the trajectory from health, through infection and sickness and back to recovery or death [5]. In particular, disease space with looping behaviour and disease maps that are circular can be used to, for example: 1.) describe the in-host dynamics of infections; 2.) identify where data of patients lie along the infection timeline, regardless of whether the stage of the infection is known (for example, from cross-sectional studies or from data of patients coming clinics to be treated, at different (and unknown) stages of the infection); and 3.) distinguish between more and less resilient individuals, from a relatively early stage of the infection course. This methodology was applied in [5] to investigate malaria. Motivated from this approach, we investigated if looping curves are a common motif in influenza or influenza in co-infection with bacteria. To that end, we also generated phase plots of pairs of features of our data sets and used K-nearest neighbour analysis, as is explained below.

##### 2.1.1 Possible looping revealed by disease maps

To help us see whether looping curves are a common motif in influenza, pneumococcal disease, or co-infection, and to help us identify those pairs of features in each of the infection groups that would be useful disease maps, we plotted the mean value at each time point of all our features (viral load, bacterial burden and cytokine concentration) pairwise against each other, for each infection group. As an example, S1 Fig. 4 shows these plots for the IAV + T4 infection group only; the other infection groups are not shown.

First, we found that the pairs in shifted phase create open loops, and not fully closed loops as is the case in [5]. Fully closed loops are not present in our data sets because data corresponds to an incomplete infection course in both the viral and the bacterial infections: The first data points in the three data sets correspond to 1.5 hours post the onset of the bacterial infection (which is the equivalent of 7 days and 1.5 hours post onset of the viral infection). Furthermore, according the ethical standards, the mice were euthanized at 31 hours post onset of bacterial infection (which is the equivalent of 8 days and 7 hours post onset of the viral infection), before the bacterial infection could resolve naturally. Therefore neither the start nor the end of the viral infection course are present and only the start of the bacterial infection course is present in the data sets. Nevertheless, to continue the search for looping behaviour and the disease maps, we closed the half loops by connecting the end points with a straight line, and choose the pairs of features that generated non-self-intersecting closed loop. Finally, following [5] we calculated the area of the resulting polygon, to make a choice of the pair of features that are "best suited" to be considered disease maps; for example, choosing those with the largest area would provide a visually clearer disease map. For the IAV + T4 infection group, the disease maps are shown in S1 Fig. 4 inside the orange boxes.

Next, we found that not the same pair of features generates a disease map for all infection groups. This means that it is not possible to use the disease maps as a tool to make direct comparisons between the behaviour of the immune system of infected individuals in the three infection scenarios (single virus, single bacterial, co-infection between the two). Notwithstanding, the disease maps can be used to investigate the individual infection groups and there is indeed an indication of possible looping motifs in the three infection groups. Therefore, to further investigate this we employed the K-nearest neighbors analysis.

##### 2.1.2 K-nearest neighbour analysis

We envisioned that if there is a true global looping structure to the data sets of each infection group, K-nearest neighbor analysis would reveal graphs that globally trace these loops and that locally show points nearby in time are connected to each other. To examine the data using nearest neighbor analysis, we took each infection group separately and stripped the data of time information. We then connected individual data points to their three nearest neighbours. Nearest neighbour analysis was performed with the `KNeighborsClassifier` function from scikit-learn [7] using the cosine distance as the metric (see section 1.1.8 for a discussion on the choice of metric). Three nearest neighbors were used as in the experimental procedure five mice were tested at each time point. The resulting k-nearest neighbour networks for each infection group showed no global looping or half-looping structures in any of the three infection groups (images not shown).

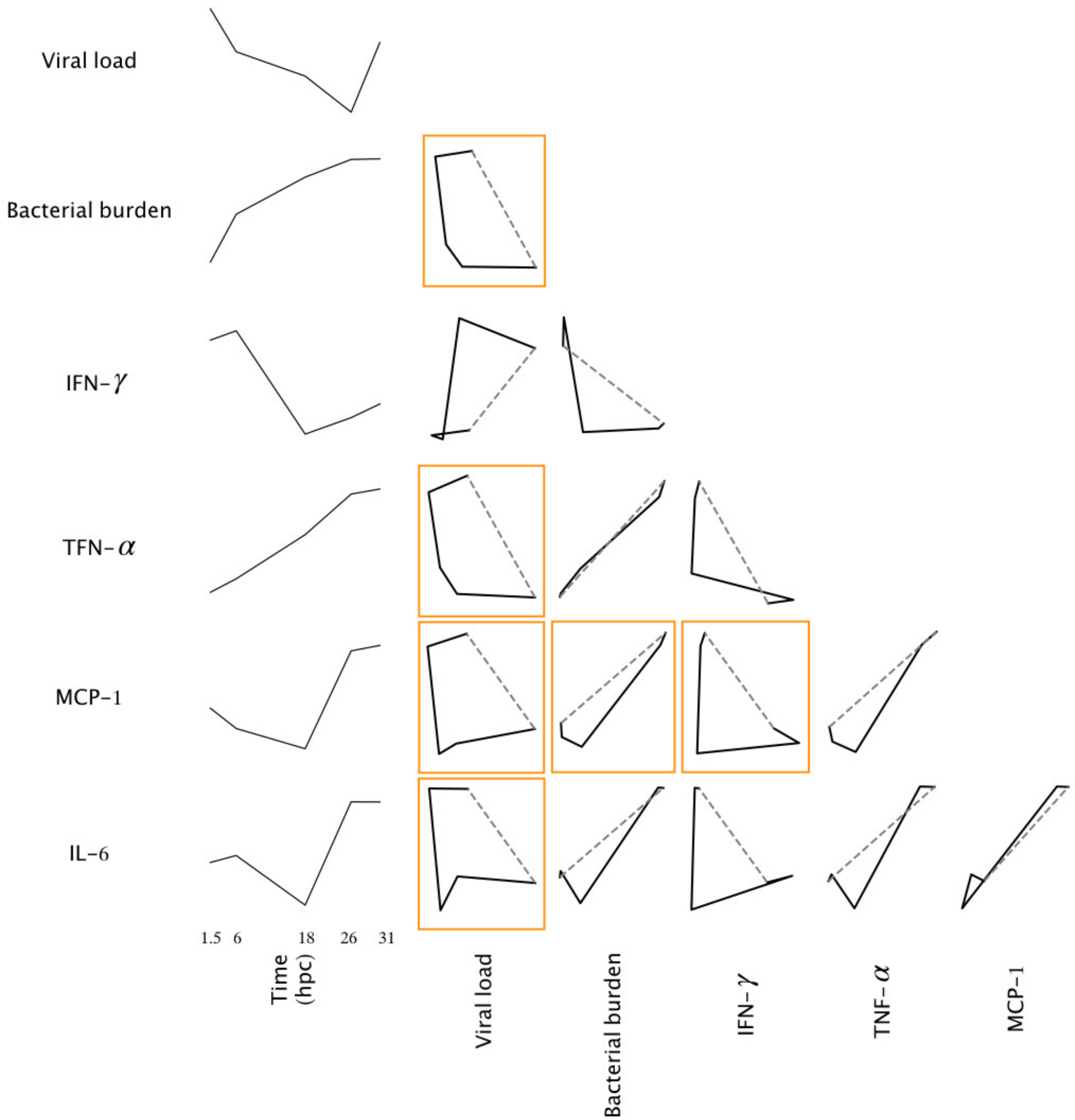


**S1 Fig. 4. Disease maps of mice in the IAV + T4 infection group.** On the first column from left to right are the time course plots of the mean values at each time point of the parasites load and the 4 core cytokines identified in [8]. The other columns are the phase space plots of the mean values at each time point for the pairs of features. Note that none of the phase plots form a completely closed loop because the data set does not have data for the complete viral nor bacterial infection courses. Therefore the loops were closed with a straight dashed line that connected the mean values of the first and last time points. Plots inside orange boxes correspond to the *disease maps* for the IAV + T4 infection group (i.e. pairs of features that generate closed non-self-intersecting curves). The phase plots not highlighted by the orange boxes are pairs of features that do not form a disease map.

#### 2.2 Parameter values for the topological networks discussed in the main script

S1 Table 1 shows the parameter values used with the Kepler mapper to generate the simplicial complexes of Figures 2 and 3 in the main script.

#### 2.3 Coloring simplicial complexes according to feature value

A strategy for evaluating the results that can be concluded from the simplicial complexes constructed by the mapper algorithm is to color the vertices according to the values of the features of the data set. More precisely, a vertex of a simplicial complexes is colored according to the average value of the feature for the data points that belong to that particular cluster. S1 Fig. 5 shows these colorings for one of the simplicial complexes for the IAV + T4 infection group. We make the following observations:

1. For the coloring corresponding to the viral load, the simplex shows high values of virus at early time points.
2. For the coloring corresponding to the bacterial burden, the simplex shows high values of bacteria at late time points.
3. For the coloring corresponding to the concentration of IFN-$\gamma$, the simplex shows high values at early time points.
4. The coloring corresponding to the concentration of TNF-$\alpha$ does not reveal anything in particular.
5. The coloring corresponding to the concentration of MCP-1 shows high values of MCP-1 at late time points.
6. The coloring corresponding to the concentration of IL-6 shows high values of IL-6 for late time points.

Note that these observations can also be made from the time course plots in S1 Fig. 4.

#### 2.4 Box plots of data points that belong to the two distinct regions revealed by TDA

In Figure 2 in the main text, the simplicial complex generated with lens 2 = IL-6 there are two regions - one in yellow (late time points IAV + T4; denoted by $g1$ in S1 Fig. 6) and one in teal/purple (early time points IAV + T4 and all time points for IAV and for T4; denoted by $g2$ in S1 Fig. 6). S1 Fig. 6 shows box plots of data points that belong to the two regions revealed by the simplicial complex. From this we observe:

1. The box plot indicates that data points of IAV + T4 from 26 and 31 hours post co-infection are separated by the simplicial complex from the earlier data points.
2. 18 and 26 hpc seem to be transition points for the system, since at both time points there are data points in both groups (the yellow and the teal/purple).

**S1 Table. 1.** Parameter values for the Kepler mapper to generate the simplicial complexes of Figures 2 and 3 in the main script. The metric used was the correlation.

| Lens 1 | Lens 2 | Num. intervals Lens 1 | Num. intervals Lens 1 | Percentage overlap Lens 1 | Percentage overlap Lens 2 | Clusterer eps |
|---|---|---|---|---|---|---|
| All infection groups | | | | | | |
| Dist. 1st Neighbour | Projection viral load lung | 1 | 13 | 0.3 | 0.8 | 0.5 |
| Dist. 1st Neighbour | Projection bacterial burden lung | 3 | 7 | 0.8 | 0.7 | 0.5 |
| Dist. 1st Neighbour | Projection IFN-$\gamma$ | 5 | 3 | 0.8 | 0.8 | 0.5 |
| Dist. 1st Neighbour | Projection TNF-$\alpha$ | 1 | 9 | 0.2 | 0.8 | 0.5 |
| Dist. 1st Neighbour | Projection MCP-1 | 5 | 3 | 0.8 | 0.4 | 0.5 |
| Dist. 1st Neighbour | Projection IL-6 | 1 | 13 | 0.3 | 0.8 | 0.5 |
| IAV | | | | | | |
| Dist. 1st neighbour | Projection viral load lung | 1 | 13 | 0.2 | 0.8 | 0.5 |
| Dist. 1st neighbour | Projection IFN-$\gamma$ | 3 | 3 | 0.6 | 0.8 | 0.5 |
| Dist. 1st neighbour | Projection TNF-$\alpha$ | 3 | 3 | 0.8 | 0.7 | 0.5 |
| Dist. 1st neighbour | Projection MCP-1 | 1 | 9 | 0.2 | 0.8 | 0.5 |
| Dist. 1st neighbour | Projection IL-6 | 3 | 3 | 0.6 | 0.8 | 0.5 |
| T4 | | | | | | |
| Dist. 1st neighbour | Projection bacterial burden lung | 3 | 3 | 0.7 | 0.8 | 1.1 |
| Dist. 1st neighbour | Projection IFN-$\gamma$ | 3 | 3 | 0.7 | 0.8 | 1.1 |
| Dist. 1st neighbour | Projection TNF-$\alpha$ | 1 | 13 | 0.2 | 0.8 | 1.1 |
| Dist. 1st neighbour | Projection MCP-1 | 3 | 3 | 0.7 | 0.8 | 1.1 |
| Dist. 1st neighbour | Projection IL-6 | 3 | 3 | 0.7 | 0.8 | 1.1 |
| IAV + T4 | | | | | | |
| Dist. 1st neighbour | Projection viral load lung | 1 | 13 | 0.3 | 0.8 | 0.5 |
| Dist. 1st neighbour | Projection bacterial burden lung | 3 | 7 | 0.7 | 0.7 | 0.5 |
| Dist. 1st neighbour | Projection IFN-$\gamma$ | 3 | 3 | 0.8 | 0.7 | 0.5 |
| Dist. 1st neighbour | Projection TNF-$\alpha$ | 5 | 3 | 0.8 | 0.6 | 0.5 |
| Dist. 1st neighbour | Projection MCP-1 | 3 | 3 | 0.8 | 0.4 | 0.5 |
| Dist. 1st neighbour | Projection IL-6 | 3 | 3 | 0.8 | 0.5 | 0.7 |

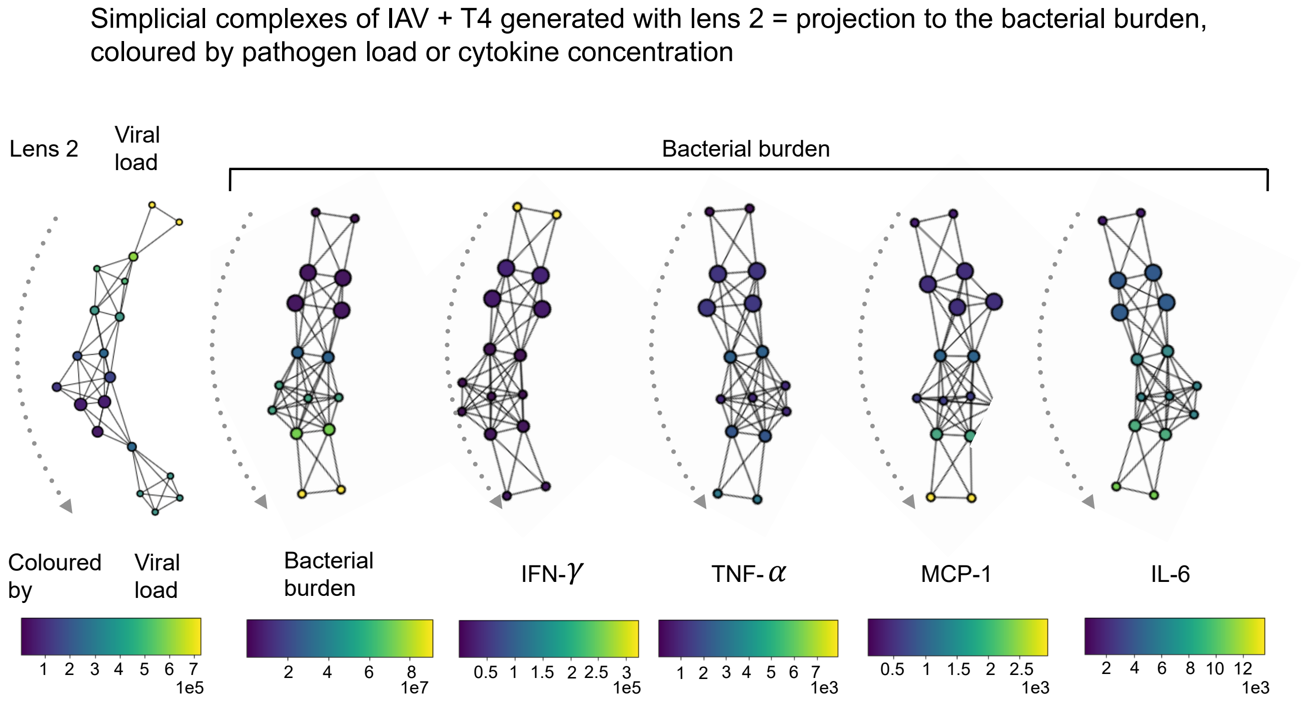


**S1 Fig. 5. Simplicial complexes of the co-infection group (IAV + T4) generated with lens 2 = projection to the viral load or bacterial burden, coloured by parasite load or cytokine concentration.** A vertex of a simplicial complex is colored according to the average value of the feature for the data points that belong to that particular cluster. The legends indicate the feature and the color maps indicate the feature values.

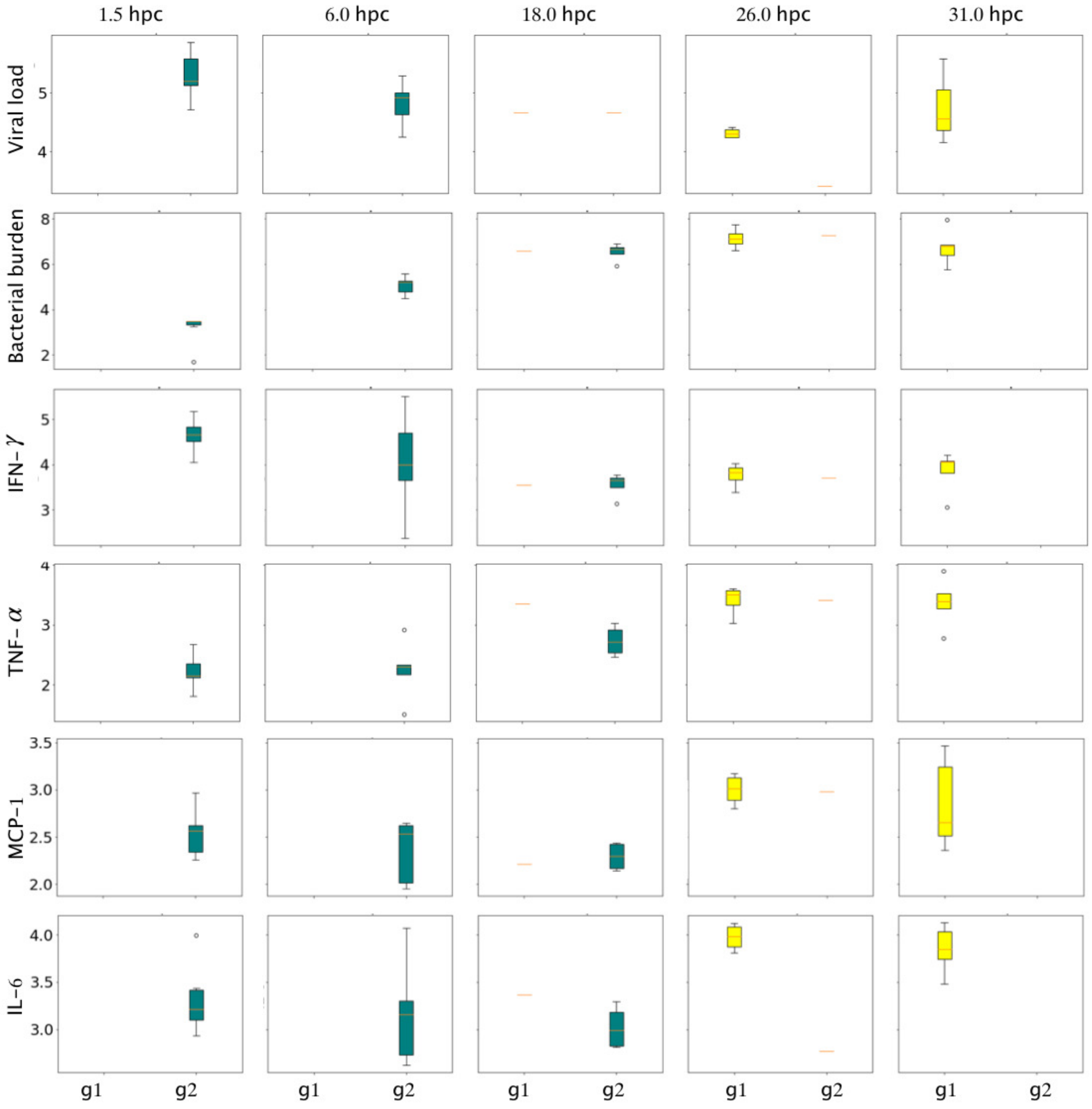


**S1 Fig. 6. Box plots of data points that belong to the two distinct regions revealed by TDA.** $g1$ denotes the data points belonging to vertices in the yellow group in Fig. 2 in the main text. $g2$ denotes the data points belonging to vertices in the teal/purple group in Fig. 2 in the main text. 6).
